# CD11b suppresses TLR7-driven inflammatory signaling to protect against lupus nephritis

**DOI:** 10.1101/2024.07.26.605143

**Authors:** Xiaobo Li, Veronica Villanueva, Viviana Jimenez, Billy Nguyen, Nishant Ranjan Chauhan, Samia Q. Khan, Jessica M. Dorschner, Mark A. Jensen, Khulood Alzahrani, Huiting Wei, David J. Cimbaluk, David C. Wei, Meenakshi Jolly, Darlah Lopez-Rodriguez, Santiago Balza Pineda, Antonio Barbosa, Roberto I. Vazquez-Padron, Hafeez M. Faridi, Jochen Reiser, Timothy B. Niewold, Vineet Gupta

**Author notes:** These authors contributed equally to this manuscript (Xiaobo Li, Veronica Villanueva, Viviana Jimenez, Billy Nguyen, Nishant Ranjan Chauhan). Current address: Department of Surgery, Division of Trauma and Critical Care, Northwestern University Chicago, IL 60612, United States. Current address: AstraZeneca, Cambridge, MA, United States. Current address: Department of Pathology, The First Affiliated Hospital Sun Yat-sen University, Guangzhou, 510060, China (Huiting Wei). Correspondence to: Vineet Gupta, PhD, Medical Research Building, Suite 9.104, 224 11^th^ Street, The University of Texas Medical Branch, Galveston, TX 77555.

## Abstract

Lupus Nephritis (LN) is a severe complication of systemic lupus erythematosus (SLE) that affects kidney function. Here, we investigated the role of CD11b, a protein encoded by the *ITGAM* gene, in the development of LN and its functional activation as a therapeutic strategy. Genetic coding variants of *ITGAM* significantly increase the risk for SLE and LN by producing a less active CD11b and leading to elevated levels of type I interferon (IFN I). However, a molecular mechanism for how these variants increase LN risk has been unclear. Here, we determined that these variants also significantly associate with elevations in soluble urokinase plasminogen activator receptor (suPAR), a known biomarker linked to kidney disease, suggesting a novel molecular connection. Pharmacologic activation of CD11b with a novel, clinical-stage agonist ONT01 significantly suppressed suPAR production in myeloid cells and reduced systemic inflammation and kidney damage in multiple experimental models of LN. Importantly, delaying treatment with ONT01 until after disease onset also significantly reduced serum suPAR and inflammatory cytokines, and decreased immune complex deposition in the glomerulus, glomerulonephritis and albuminuria, suggesting that CD11b activation is therapeutic for LN. Genetic activation of CD11b via a gain-of-function CD11b mutation also showed complete protection from LN, whereas genetic deletion of CD11b worsened the disease in mice, providing further evidence of the role of CD11b activation in regulating LN. Finally, transfer of human LN PBMCs generated human LN like disease in mice that was significantly reduced by ONT01. Together, these data provide strong evidence that ONT01 mediated CD11b activation can therapeutically modulate TLR7-driven inflammation and protect against LN. These findings support clinical development of CD11b agonists as novel therapeutics for treating lupus nephritis in human patients.

## Introduction

Systemic lupus erythematosus (SLE) is a highly heterogenous autoimmune disease with dysregulated innate and the adaptive immune responses that result in tissue damage and multi-organ dysfunction [1–3]. Over half of all SLE patients develop lupus nephritis (LN), an inflammatory kidney damage that is the strongest predictor of morbidity and mortality in patients [4, 5]. Yet, the precise molecular mechanisms driving SLE and its progression to LN remain unclear, hampering development of novel and targeted therapeutics.

Toll-like receptors (TLRs) play a key in the pathogenesis of SLE and LN. TLRs are pattern recognition receptors that detect pathogen-associated molecular patterns (PAMPs) and damage-associated molecular patterns (DAMPs), such as single-stranded RNA (ssRNA) and double-stranded DNA (dsDNA), triggering innate immune activation. Excessive TLR activation initiates downstream signaling to produce soluble urokinase plasminogen activator receptor (suPAR), a known risk factor for glomerular diseases [6–9], type I interferons (IFN I) that are heritable and pathogenic risk factors for SLE and LN [10, 11]. It also increases immune complex formation that get deposited in the glomeruli thereby increasing pro-inflammatory signaling in various kidney cells. These deposits also recruit inflammatory immune cells from circulation that increase local cytokine levels and further contribute to loss of glomerular integrity and kidney function [12, 13]. Among the various TLRs, Toll-like receptor 7 (TLR7) is a key receptor that is expressed in B cells and myeloid cells and is involved in the development and progression of SLE and LN [14–19]. Single nucleotide polymorphisms (SNPs) in the TLR7 gene result in the overexpression and hyperactivation of TLR7 that amplifies the immune response to self-antigens and are associated with increased risk of SLE and LN [20]. TLR7 detects ssRNA that triggers a signaling cascade that activates transcription factors, such as NF-κB and interferon regulatory factors 3 and 7 (IRF3 and IRF7), and induce production of pro-inflammatory cytokines and IFN I thereby exacerbating kidney injury, tubulointerstitial damage and glomerulosclerosis [1, 21–23].

Integrin CD11b, α-subunit of the heterodimeric receptor CD11b/CD18 (also known as αMβ2, CR3, Mac-1), also plays a key role in modulating immune responses. CD11b is predominantly expressed on myeloid cells, including macrophages and neutrophils, and is involved in processes such as cell adhesion, migration, phagocytosis, and the regulation of inflammatory responses [24, 25]. Genome wide associate studies (GWAS) identified SNPs in the coding region of the *ITGAM* gene, that encodes CD11b, that significantly increase the risk and are causal missense variants for SLE, LN, Sjogren’s syndrome, systemic sclerosis and other related autoimmune diseases [26–31]. Furthermore, these CD11b variants impair CD11b’s ability to bind ligands and clear immune complexes and apoptotic cells, without affecting its expression levels, thereby exacerbating the inflammatory milieu that is characteristic of SLE and LN [32–38].

suPAR is a soluble form of the urokinase plasminogen activator receptor (uPAR) that is expressed on various cell types, including myeloid cells [6–9, 39, 40]. We and others have previously shown that suPAR levels are significantly elevated in various kidney diseases, including LN, and correlate with disease severity and progression [7–9, 39]. Moreover, suPAR elicits its pathogenic effects on kidney function, by inducing podocyte injury, proteinuria, and glomerulosclerosis, through binding to integrin αVβ3 on podocytes [8].

Recently, we and others discovered that CD11b acts as a negative regulator of the pro-inflammatory TLR4 signaling pathways in SLE [32, 33, 41, 42]. We found that *ITGAM* coding SNPs that result in less-active CD11b correlate with high IFN I levels in the sera of SLE patients [41]. Inactivity or absence of CD11b failed to suppress TLR4-mediated pro-inflammatory signaling in cells, suggesting a mechanistic basis for the association between mutant CD11b and SLE. We also determined that allosteric activation of CD11b rescued the functional deficit in CD11b, *in vitro* and *in vivo*, and reduced SLE in murine models [41], suggesting that CD11b activation is a novel therapeutic mechanism for inflammatory diseases.

However, several questions remain. *First*, the mechanistic link between *ITGAM* SNPs and LN, and how these variants lead to the increased inflammation and kidney damage, is unclear. *Second*, it is not fully understood if CD11b functionally regulates TLR7, the main TLR implicated in SLE and LN. *Third*, it remains to be established whether activation of integrin CD11b can therapeutically treat TLR7-dependent autoimmunity in mice and whether this mechanism has potential for clinical development in LN. *Finally*, it is untested whether human PBMCs can induce LN in murine models and whether treatment with CD11b activators can suppress it, thereby helping translate this mechanism into therapeutics for patients. Addressing these questions could provide crucial molecular insights into LN and a novel therapeutic mechanism that could serve as a foundation for future clinical trials and the development of an innovative innate immune cell targeting therapy for this unmet medical need.

Here, we present data addressing these key questions. We demonstrate that *ITGAM* SNPs correlate LN and with increased levels of suPAR, a specific kidney disease driver. Using a TLR7-dependent LN model [43], we show that pharmacologic activation of CD11b with ONT01, an oral CD11b agonist, effectively treats LN *in vivo*. Furthermore, using a novel CD11b knock-in mouse model expressing a gain-of-function active mutant of CD11b that mimics pharmacologic activation of CD11b [44], we observe robust protection against TLR7-dependent LN. This suggests that CD11b activation serves as a therapeutic mechanism to suppress TLR7-dependent autoimmune disease. Our studies also show that transferring LN PBMCs recapitulates human LN in mice and that treatment with the CD11b agonist ONT01 significantly reduces disease severity. Collectively, our findings identify CD11b as a key regulator of TLR7-dependent inflammatory diseases, demonstrate that CD11b activation is protective against autoimmune kidney disease that support the development of CD11b agonists as potential therapeutics for LN and other autoimmune diseases.

## Results

### LN subjects carrying *ITGAM* SNPs show significant elevation in their serum suPAR levels

Perturbation and over-activation of the innate immune system results in increased serum levels of type I interferon (IFN-I) that are associated with LN [45]. Three *ITGAM* coding SNPs result in overactive myeloid cells and strongly associate with risk for SLE and LN [26–28, 46–49]. These SNPs produce less active integrin CD11b protein product that correlates with significantly elevated serum IFN-I levels in SLE subjects [41]. We and others recently showed that increased serum levels of soluble urokinase plasminogen activator receptor (suPAR) strongly associate with acute and chronic kidney diseases and strongly predict decline in kidney function [7–9, 39, 40]. Given the association of *ITGAM* SNPs with overactive myeloid cells [7], we hypothesized that these *ITGAM* SNPs mediate kidney dysfunction by increasing serum suPAR levels in LN. To test, *first*, we quantified suPAR levels in the sera of LN patients and determined that they are significantly increased as compared to the levels in healthy sera (**Fig. 1A**), consistent with findings from other recent publications [50, 51]. Next, we quantified suPAR levels in SLE patient sera. Our data shows that serum suPAR levels were significantly increased in SLE patients with renal dysfunction as compared to patients without renal function decline (**Fig. 1B**), further confirming the link between increased serum suPAR levels in chronic kidney disease [7, 39]. Similarly, SLE patients with proteinuria showed significantly elevated serum suPAR levels as compared to patients with normal levels (**Fig. 1C**). To directly link *ITGAM* SNPs with serum suPAR levels and SLE patients with renal dysfunction, we quantified suPAR in *ITGAM* SNP-carrying patients. Data shows significantly higher levels of serum suPAR among the patients carrying *ITGAM* rs1143679 risk allele and from European ancestry as compared with the non-risk allele carrier SLE patients with renal dysfunction (**Fig. 1D**). Together, these data suggest that increased serum suPAR is linked to kidney disease via *ITGAM* SNPs, suggesting a direct link between reduced CD11b activity, suPAR and lupus nephritis.

**Figure 1.**
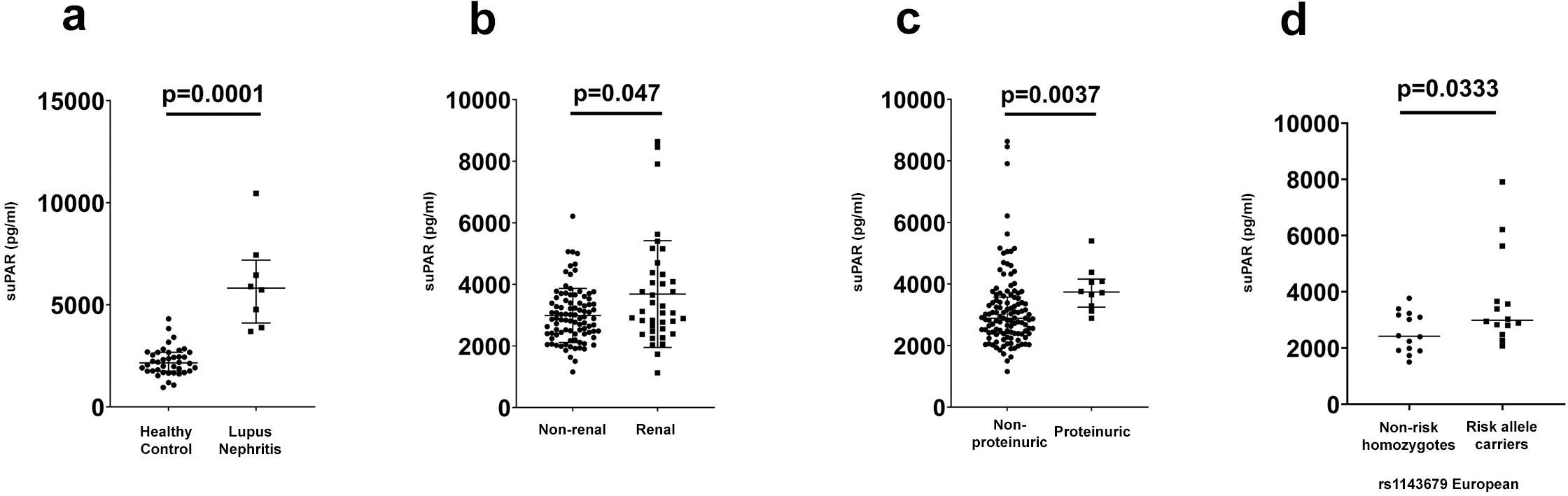
*ITGAM* SNPs are associated with elevated suPAR levels in SLE patients with renal disease. **(a)** Graph showing ELISA based quantification of suPAR levels in the sera of healthy control and lupus nephritis patients. Each dot represents a unique sample. Data show mean ± SEM. ****p < 0.0001. **(b)** Graph showing ELISA based quantification of suPAR levels in the sera of SLE patients stratified by kidney function non-renal SLE versus renal dysfunction) Each dot represents a unique sample. Data show mean ± SEM. *p < 0.05. **(c)** Graph showing ELISA based quantification of suPAR levels in the sera of SLE patients stratified by proteinuria (non proteinuric versus proteinuric). Each dot represents a unique sample. Data show mean ± SEM. **p < 0.01. **(d)** Graph showing ELISA based quantification of suPAR levels in the sera of SLE patients stratified by the *ITGAM* SNP genotype (rs1143679) non-carriers versus carriers showing correlation of *ITGAM* SNP carriers with high suPAR levels. Each dot represents a unique sample. Data show mean ± SEM. *p < 0.05.

### TLR ligands increase suPAR levels in myeloid cells and pharmacologic activation of CD11b suppresses it

Myeloid cells are the primary source of serum suPAR that results in proteinuric kidney damage [7]. Activation of TLRs on immune cells increases the production of IFN-I and other proinflammatory mediators, playing a crucial role in SLE and LN pathobiology in both murine models and humans [11, 14–20, 41, 52, 53]. To determine if TLR activation of myeloid cells also increases suPAR production, we stimulated murine RAW 264.7 macrophage cell line with macrophage activator phorbol 12-myristate-13-acetate (PMA) or with various TLR agonists (TLRa) and quantified suPAR levels in cell supernatants. Treatment with PMA, TLR4 agonist lipopolysaccharide (LPS), TLR2 agonists Pam2CSK4 (Pam2) or Pam3CSK4 (Pam3), TLR3 agonist poly I:C, TLR9 agonist CpG or TLR7 agonist imiquimod (IMQ) resulted in significantly increased levels of suPAR in cell supernatants (**Fig. 2a**). We previously showed that CD11b activation with the allosteric activator ONT01 (previously GB1275 and referred to here as ONT01 as its current clinical nomenclature [41, 54–58]) suppresses LPS-mediated TLR4-dependent pro-inflammatory signaling in myeloid cells [41]. ONT01 binds to the ligand binding α/I domain of CD11b, stabilizing its active conformation, and thereby promoting CD11b activation [58]. To test if ONT01 mediated CD11b activation suppresses other TLR-dependent increases in suPAR, we co-treated cells with TLRa and ONT01, and found that ONT01 significantly reduced suPAR levels that suggests that CD11b activation suppresses TLRa-induced suPAR production (**Fig. 2a**). TLR7 ligand induced over-activation is a known driver of SLE and blocking TLR7 signaling has therapeutic potential [20, 59]. Thus, to further explore if CD11b can suppress TLR7 signaling activity, we treated these cells with TLR7 ligand imiquimod (IMQ) (**Fig. 2b**) and quantified pro-inflammatory mediators suPAR, IFN-β, IL-6, and TNF-α in cell supernatants. We found that TLR7 ligand IMQ significantly increased secretion of these cytokines, similar to TLR4 ligand LPS [41], and that co-treatment with ONT01 significantly suppressed their increase. Quantification of uPAR, IFN-β, IL-6, and TNF-α mRNA using qPCR showed that TLR ligands transcriptionally increased these inflammatory mediators and that ONT01 significantly reduced their mRNA levels, suggesting that the effect of ONT01 is mediated at a transcriptional level (**Figs. 2c**). Blocking NF-κB, a known transducer of TLR-dependent inflammatory signaling with a known NF-κB antagonist BMS345541 [60], reduced uPAR mRNA levels, suggesting that uPAR and suPAR production is NF-κB dependent (**Fig. S1**).

**Figure 2.**
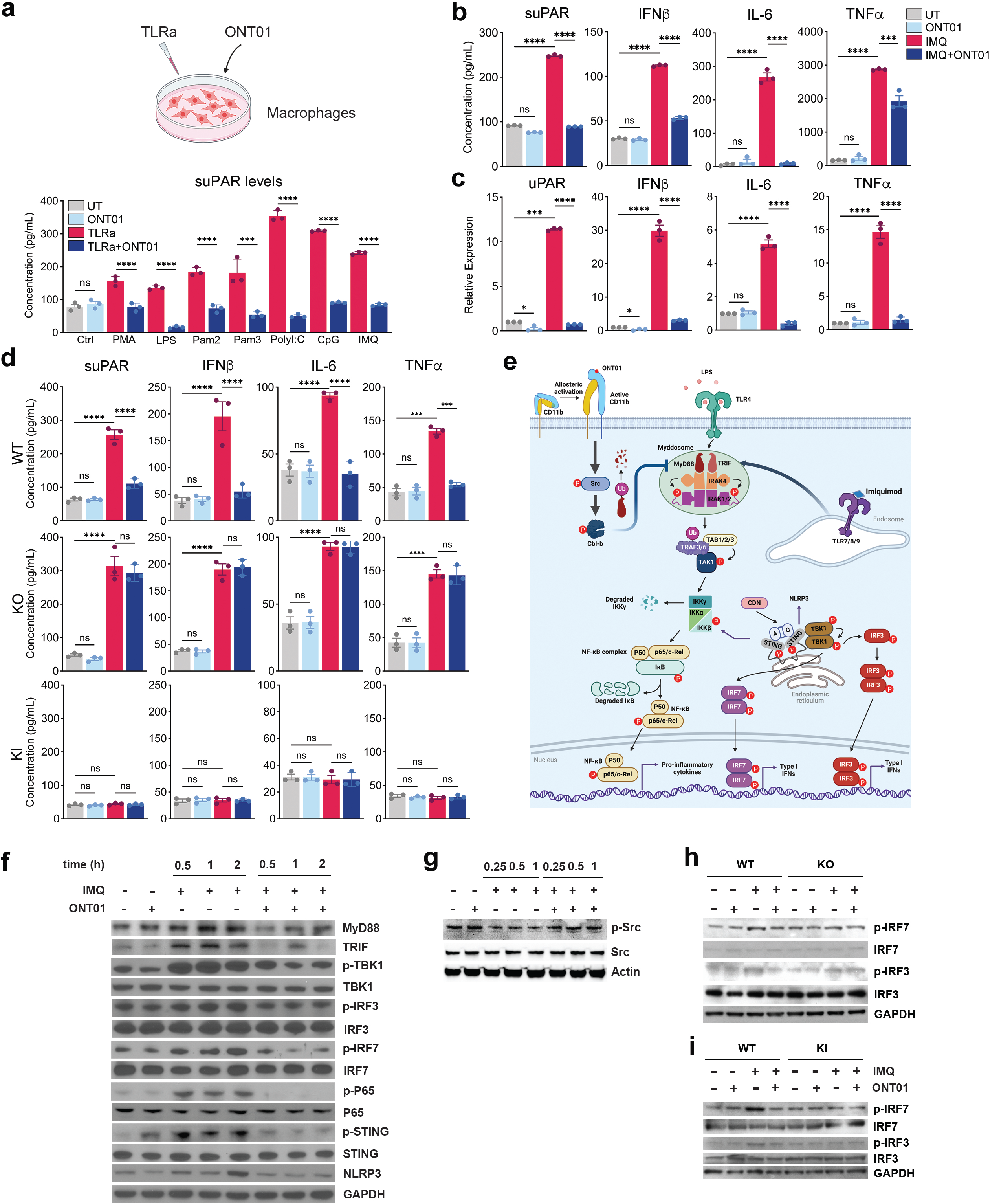
Pharmacological activation of CD11b suppresses TLR mediated increase suPAR levels in myeloid cells. **(a)** (Top) Experimental diagram showing plan of *in vitro* treatment of macrophages and cell line with TLR agonists (TLRa) and CD11b activator ONT01. (Bottom) Graph showing ELISA based quantification of suPAR levels in the supernatant of RAW 264.7 macrophage cell line cells after various treatments: control (Ctrl) untreated (UT) (gray bars) or treated with ONT01 alone (light blue bars), cells stimulated with phorbol 12-myristate-13-acetate (PMA) or with various TLR ligands/agonists (TLRa) including TLR4 agonist lipopolysaccharide (LPS), TLR2 agonist Pam2CSK4 (Pam2) or Pam3CSK4 (Pam3), TLR3 agonist poly I:C, TLR9 agonist CpG and TLR7 agonist imiquimod (IMQ) in the absence (red bars) or presence of ONT01 (blue bars). Each bar shows mean ± SEM (n=3). ns = not significant, ***p<0.001, ****p<0.0001. **(b)** Graph showing ELISA based quantification of suPAR, IFN-β, IL-6 and TNF-α levels in the RAW 264.7 macrophage cell supernatant from the following groups: control untreated cells (gray), cells treated with ONT01 alone (light blue), or cells stimulated with TLR7a IMQ (20 μg/ml) and in the absence (red bars) or presence of ONT01 (20 μM) (blue bars). Each bar shows mean ± SEM (n=3). ns = not significant, *p<0.05, ***p<0.001, ****p<0.0001. **(c)** Graph showing quantification of mRNA expression levels of uPAR, IFN-β, IL-6 and TNF-α by qRT-PCR levels in the RAW 264.7 macrophages, as in (**b**). Each bar shows mean ± SEM (n=3). ns = not significant, *p<0.05, ***p<0.001, ****p<0.0001. **(d)** Graph showing ELISA based quantification of suPAR, IFN-β, IL-6 and TNF-α levels in cell supernatants of BMDMs from WT, KO, and KI mice. Data shows measurements from control untreated cells (UT, gray bars), cells treated with ONT01 alone (ONT01, light blue bars), or from cells stimulated with TLR7a IMQ (20 μg/ml) and in the absence (TLR7a, red bars) or presence of ONT01 (20 μM) (TLR7a + ONT01, blue bars). Each bar shows mean ± SEM (n=3). ns = not significant, *p<0.05, ***p<0.001, ****p<0.0001. **(e)** A schematic showing regulation of various molecular components of the TLR7/TLR4 mediated type I IFN and NF-kB signaling pathways upon CD11b activation (with ONT01). **(f)** Images showing immunoblot analysis of various phosphorylated proteins (p-TBK1, p-IRF3, p-IRF7, p-p65, p-STING) and total proteins in the TLR7 signaling pathway in lysates of RAW macrophages stimulated with IMQ (20 μg/ml) in the absence or presence of ONT01 (20 μM) for 0.5 h, 1 h and 2 h. GAPDH was used as loading control. **(g)** Images showing immunoblot analysis of phosphorylated Src (pSrc) and total Src in the TLR7 signaling pathway in lysates of RAW macrophages stimulated with IMQ (20 μg/ml) in the absence or presence of ONT01 (20 μM) for 0.25 h, 0.5 h and 1 h. Actin was used as loading control. **(h)** Images showing immunoblot analysis of p-IRF7, p-IRF3, total IRF7 and IRF3 in lysates of BMDMs from WT, KO and KI mice and stimulated with IMQ (20 μg/ml) in the absence or presence of ONT01 (20 μM) for 1 h. GAPDH was used as loading control.

### CD11b deficiency increases and CD11b activation suppresses TLR7-stimulation dependent suPAR levels in primary myeloid cells

To examine CD11b’s regulatory role in TLR7-dependent pro-inflammatory pathways, we isolated bone marrow cells from wild-type C57BL/6 (WT), CD11b knockout (CD11b KO), and transgenic CD11b knock-in (CD11b KI) mice with a gain-of-function I332G mutation that renders CD11b constitutively active [44, 61]. Cultured bone marrow-derived macrophages (BMDMs) from WT mice showed increased suPAR, IFN-β, IL-6, and TNF-α levels upon IMQ stimulation, which were significantly reduced by ONT01 (**Fig. 2d**, WT panel). CD11b KO BMDMs produced higher levels of these markers with IMQ, indicating unrestrained TLR7 signaling in the absence of CD11b (**Fig. 2d**, KO panel). ONT01 did not reduce these markers in KO cells, confirming its CD11b-dependent function [41, 42, 56–58, 62]. CD11b KI cells showed no significant increase in suPAR, IFN-β, IL-6, and TNF-α levels with IMQ, mirroring the pharmacologic effects of ONT01 (**Fig. 2d**, KI panel). Co-treatment of KI cells with ONT01 showed no additional decrease, suggesting optimal CD11b activity in KI cells. The mRNA quantification mimicked these results (**Fig. S2**), suggesting that the observed changes in in suPAR, IFN-β, IL-6, and TNF-α are transcriptionally regulated. These findings were also consistent with TLR4 stimulation (**Fig. S3**). Thus, both genetic and pharmacologic CD11b activation suppress TLR7-dependent increases in suPAR and other pro-inflammatory markers.

### CD11b activation suppresses TLR7-dependent NF-κB proinflammatory pathways

To examine how CD11b activation protects against TLR7-dependent pro-inflammatory signaling, we performed western blot (WB). TLRs activate intracellular inflammatory signaling by binding various ligands and assembling adaptors MyD88 and TRIF into dynamic, barrel like structures called myddosomes [63]. Myddosomes soon dissociate from the membrane associated TLRs and organize effector proteins, including adaptors and kinases. We previously showed that CD11b activation suppresses TLR4-dependent NF-kB and IRF3/7 pathways [41]. WB analyses showed that CD11b activation down-modulated TLR7a IMQ-mediated increase in MyD88 and TRIF by inducing Src phosphorylation [32] (**Figs. 2e-2g**). ONT01 normalized the IMQ-induced increase in phospho-p65 (p-p65), down-modulating the TLR7-dependent pro-inflammatory NFκB pathway. WB analyses also showed ONT01 reduced IMQ-mediated increases in pIRF3 and pIRF7 levels. TANK-binding kinase 1 (TBK1) phosphorylation of IRF3 and IRF7 is crucial for IFN I production [64, 65]. Phospho-TBK1 activates the pro-inflammatory Stimulator of Interferon Genes (STING) and NLR family pyrin domain containing 3 (NLRP3) pathways [65–67]. ONT01 treatment normalized IMQ-induced increases in pTBK1, pSTING, and NLRP3 levels, indicating that CD11b activation suppresses TLR7-dependent IFN I pathways via pIRF3 and pIRF7 down-modulation.

In primary murine BMDM cells, IMQ stimulation of WT BMDMs increased pIRF3 and pIRF7 levels, which were suppressed by ONT01 (**Figs. 2g and S4**). CD11b KO BMDMs showed basal increases in these proteins, confirming the over-active IFN I pathway in CD11b KO macrophages [41]. ONT01 had no effect on KO cells (**Figs. 2g and S4a**), verifying its CD11b-mediated action. CD11b KI BMDMs showed no IMQ-induced increases in pIRF3 and pIRF7 levels, and ONT01 did not further reduce their basal levels (**Figs. 2g and S4b**), confirming genetic CD11b activation regulates TLR7-dependent signaling.

To examine if direct NF-κB activation can override the inhibitory effect of upstream CD11b activation, we treated CD11b KI BMDMs with an NF-κB or a STING agonist. This treatment increased suPAR and IL-6, indicating that direct NF-κB activation bypasses the TLR/MyD88 block induced by activated CD11b (**Fig. S5**). Collectively, these data show that pharmacologic and genetic activation of CD11b regulates TLR7-dependent pro-inflammatory signaling by down-modulating MyD88/TRIF-driven NFκB and IRF3/7 pathways in myeloid cells.

### CD11b activation therapeutically protects against TLR7-induced LN

To investigate CD11b’s protective effects in TLR7-dependent autoimmunity, we used a TLR7-induced LN model [14, 43] by treating WT animals epicutaneously with TLR7a IMQ for 8 weeks (**Fig. 3a**). IMQ-treated mice developed a systemic lupus phenotype, including splenomegaly (**Figs. 3b** and **S6**), splenic expansion of CD11b+ leukocytes (**Fig. 3c**), increased serum anti-dsDNA antibodies, suPAR, and IL-6 (**Fig. 3d**), and significant albuminuria and histologic kidney damage, suggesting systemic inflammation driven end organ disease (**Figs. 3e-i**).

**Figure 3.**
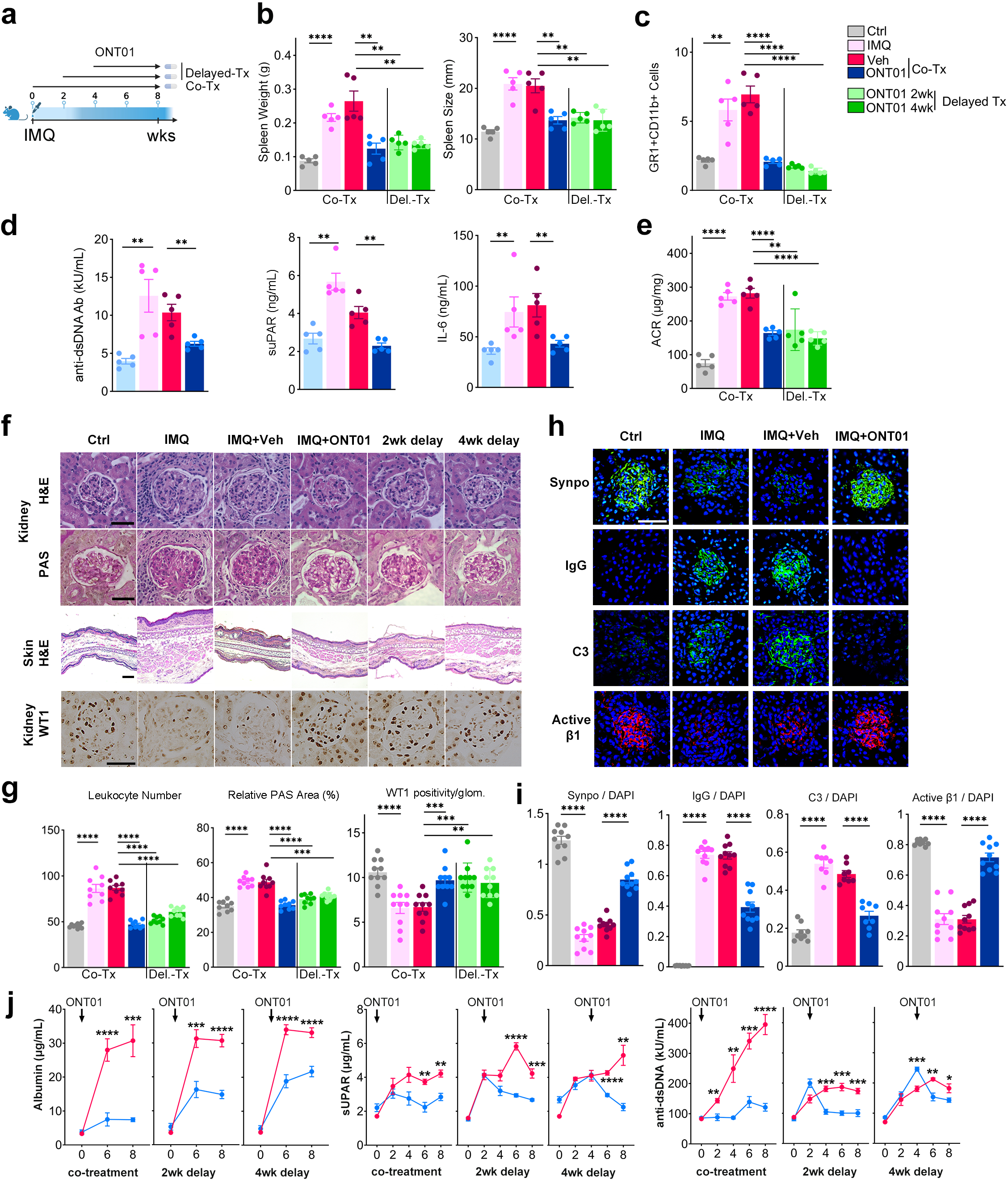
Pharmacologic activation of CD11b therapeutically protects against TLR7-induced LN. **(a)** Experimental diagram showing plan of *in vivo* treatment of mice with TLR7a IMQ and CD11b activator ONT01. Arrows indicate the duration of treatment with ONT01. Treatment was initiated either simultaneously with initiation of IMQ (co-treatment), 2 weeks after initiation of LN by IMQ (2-wk delayed) or 4 weeks after initiation of LN by IMQ (4-wk delayed). **(b)** Graphs showing quantification of spleen weight (in g) and size (in mm) at the end point from untreated animals (UT, gray bars), or treated with IMQ alone (1.25 mg of 5% cream, pink bars), with IMQ and co-treated with vehicle (red bars), with IMQ and co-treated with oral ONT01 (30 mg/kg, BID, blue bars), with IMQ and treated with ONT01 after a 2-wk delay (light green bars), or with IMQ and treated with ONT01 after a 4-wk delay (dark green bars). Each bar shows mean ± SEM (n=5 animals/group). **p<0.01, ****p<0.0001. **(c)** Graph showing flow cytometric analysis of GR1+CD11b+ splenocytes from various groups of animals, as in **b**. Each bar shows mean ± SEM (n=5 animals/group). **p<0.01, ****p<0.0001. **(d)** Graphs showing ELISA based quantification of anti-dsDNA antibody, suPAR and IL-6 levels in the end-point sera from various groups of animals, as in **b**. Each bar shows mean ± SEM (n=5 animals/group). *p<0.05, **p<0.01. **(e)** Graph showing urinary albumin to creatinine ratio (ACR) in the end-point urine from various groups of animals, as in **b**. Each bar shows mean ± SEM (n=5 animals/group). **p<0.01, ****p<0.0001. **(f)** Representative images from end point kidney (stained with H&E and PAS) and skin (stained with H&E) sections from various groups of animals, as in **b.** Also presented are representative images of kidney glomeruli from immunohistochemical staining of kidney with antibodies against Wilms Tumor 1 (WT1, brown) from various groups of animals, as in **b**. Scale bar: 50 μm for kidney; 100 μm for skin. **(g)** Bar graphs showing quantification of leukocyte counts, relative PAS area, and WT1 positive cell numbers (indicating number of podocytes) among the experimental groups presented in **f**. Each bar shows mean ± SEM (n=8-10 independent images/group). **p<0.01, ***p<0.001, ****p<0.0001. **(h)** Representative immunofluorescence images of kidney glomeruli from staining of kidney with antibodies against synaptopodin (Synpo, green), IgG (IgG, green), complement (C3, green) and integrin β1 activation epitope (9EG7, red) from animals as in **i**. Nuclei were stained with DAPI (blue). Scale bar: 50 μm. **(i)** Bar graphs showing quantified ratios of IgG and C3 deposition in glomerulus of the various experimental groups presented in **m**. Each bar shows mean ± SEM (n=8-10 independent images/group). ****p<0.0001.

Given our recent data showing CD11b activation to be protective in SLE models [41], we studied the effect of using CD11b activator in this TLR7-induced LN model. Oral administration of ONT01 significantly protected mice from systemic lupus and inflammation (dark blue bars, **Figs. 3b-3i** and **Fig. S6**). ONT01 normalized spleen size and weight, significantly reduced splenic expansion of CD11b^+^ leukocytes (**Fig. 3c**) and decreased circulating anti-dsDNA antibodies, suPAR, and IL-6 compared to vehicle treated animals (dark blue vs red bars, **Fig. 3d**). ONT01 also significantly reduced albuminuria (**Fig. 3e**), decreased glomerular damage, reduced mesangial expansion and kidney leukocyte infiltration, and reduced skin inflammation (**Fig. 3f-3g**). To validate ONT01’s protective effect on glomerular damage, we performed histologic and IF analyses on glomerular tissues. IF analysis of kidney glomeruli using an integrin β1 activation-specific antibody (9EG7) and an anti-Synpo antibody showed significant podocyte loss in IMQ and IMQ+vehicle-treated animals. This loss was significantly reduced by ONT01 (**Figs. 3h-3i**). IHC staining with the podocyte-expressed transcription factor WT1 further confirmed that ONT01 significantly reduced podocyte cell loss induced by IMQ (**Figs. 3f-3i**).

Together, these data suggest that CD11b agonist reduces TLR7-induced myeloid activation *in vivo*, reduces kidney damage biomarkers suPAR and IL-6 in circulation, reduces leukocyte infiltration and immune complex and C3 deposition in the kidney, and reduces podocyte damage and loss, to ameliorate LN.

To investigate whether CD11b activation is therapeutic, we delayed administration of ONT01 to IMQ-treated mice by 2-wks or 4-wks after disease induction, after the animals present with significant disease. These delayed ONT01 treatments also significantly reduced albuminuria and serum inflammatory biomarkers within two weeks (**Fig. 3j**). Histologic and IF analyses confirmed protection from mesangial expansion, leukocyte infiltration, and podocyte damage and loss in both delayed treatment groups (**Figs. 3f-3i**). Spleen size, weight, and splenic CD11b+ leukocyte numbers normalized in both groups (green bars, **Figs. 3b-3c**), confirming ONT01’s therapeutic efficacy.

To further establish the role of CD11b activation as protective in LN, we used transgenic mice expressing genetically activated CD11b with a gain-of-function I332G mutation (CD11b KI (KI) [44]) and compared them to WT and CD11b KO (KO) mice [68]. We treated these mice with IMQ for 8 weeks (**Fig. 4a**). CD11b KO animals, prone to autoimmune disease [69], exhibitied worsened splenomegaly and splenic CD11b^+^ leukocyte expansion compared to WT mice (**Figs. 4b-4c**). KO mice also had high serum anti-dsDNA, suPAR and IL-6 levels (**Figs. 4d-4f**), along with increased albuminuria and histologic kidney damage (**Figs. 4g-4h**), and increased renal deposition of IgG and C3 immune complexes (**Figs. 4j-4k**), indicating that CD11b is essential for restraining TLR-dependent responses and for suppressing systemic inflammation.

**Figure 4.**
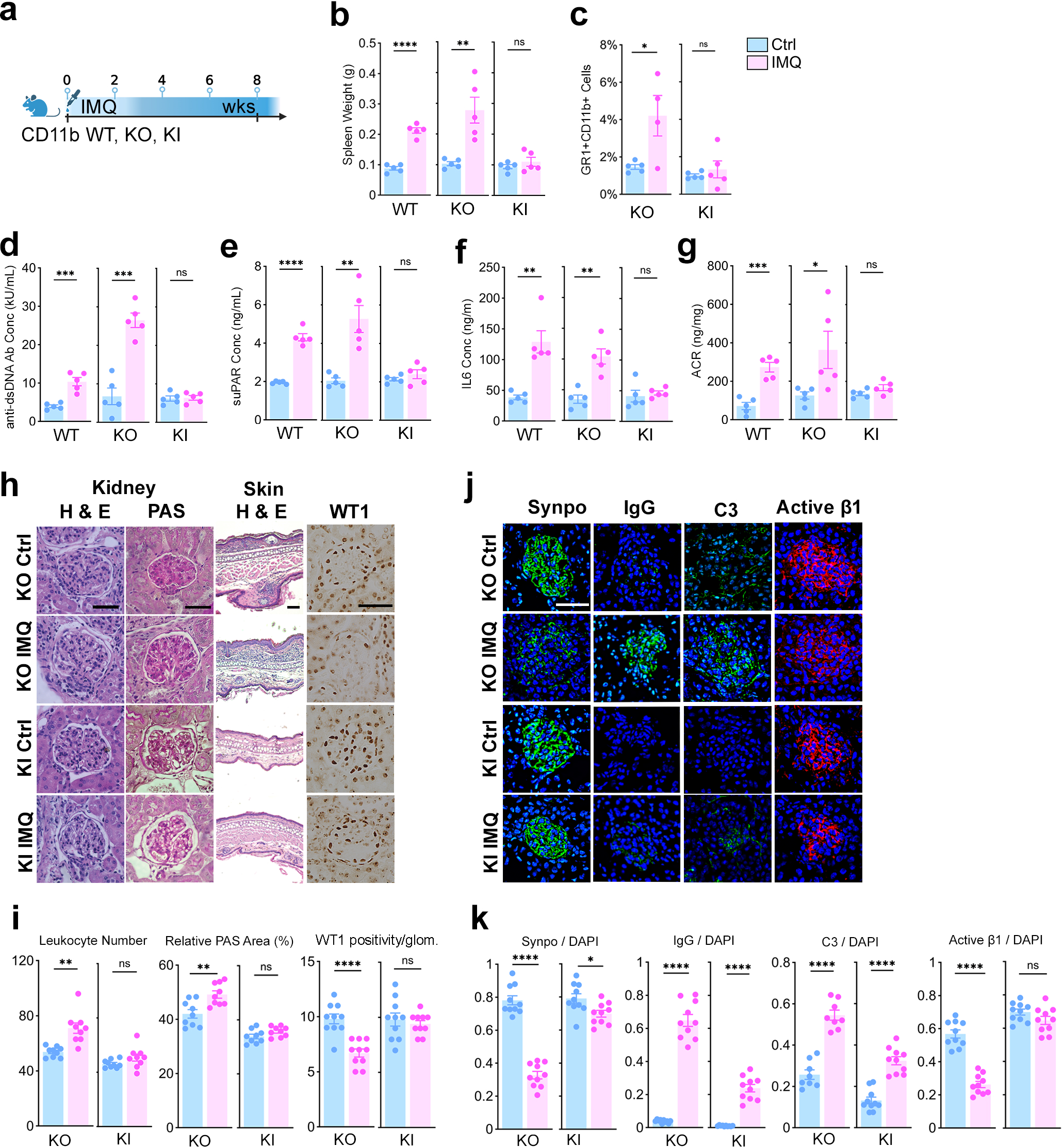
Genetic activation of CD11b protects against LN. **(a)** Experimental diagram showing plan of *in vivo* treatment of CD11b WT, KO and KI mice with TLR7a IMQ. **(b)** Graph showing quantification of spleen weight (in g) at the end point from untreated animals (UT, blue bars), or treated with IMQ (1.25 mg of 5% cream, pink bars). Each bar shows mean ± SEM (n=5 animals/group). ns=not significant, *p<0.05, **p<0.01, ****p<0.0001. **(c)** Graph showing flow cytometric quantification of GR1+CD11b+ splenocytes from various groups of animals, as in **b**. Each bar shows mean ± SEM (n=5 animals/group). **p<0.01, ****p<0.0001. **(d-f)** Graphs showing ELISA based quantification of anti-dsDNA antibody, suPAR and IL-6 levels in the end-point sera from various groups of animals, as in **b**. Each bar shows mean ± SEM (n=5 animals/group). ns=not significant, **p<0.01, ****p<0.0001. **(g)** Graph showing albumin to creatinine ratio (ACR) in the end-point urine from various groups of animals, as in **b**. Each bar shows mean ± SEM (n=5 animals/group). **p<0.01, ****p<0.0001. **(h)** Representative images from end point kidney (stained with H&E and PAS) and skin (stained with H&E) sections from various groups of animals, as in **b.** Also presented are representative images of kidney glomeruli from immunohistochemical staining of kidney with antibodies against Wilms Tumor 1 (WT1, brown) from various groups of animals, as in **b**. Scale bar: 50 μm for kidney; 100 μm for skin. **(i)** Bar graphs showing quantification of leukocyte number, relative PAS area, and WT1 positive cell numbers among the experimental groups presented in **h**. Each bar shows mean ± SEM (n=8-10 independent images/group). ns=not significant, **p<0.01, ****p<0.0001. **(j)** Representative immunofluorescence images of kidney glomeruli from staining of kidney with antibodies against synaptopodin (Synpo, green), IgG (IgG, green), complement (C3, green) and integrin β1 activation epitope (9EG7, red) from various groups of animals, as in **h**. Nuclei were stained with DAPI (blue). Scale bar: 50 μm. **(k)** Bar graphs showing quantified ratios of IgG and C3 deposition in glomerulus of the various experimental groups presented in **l**. Each bar shows mean ± SEM (n=8-10 independent images/group). ****p<0.0001.

Conversely, the KI animals expressing constitutively active CD11b showed complete protection from systemic lupus and LN, with no increase in spleen weight (**Fig. 4b**), no elevations in anti-dsDNA antibodies, suPAR or IL-6 levels, and no increase in albuminuria or histologic kidney damage (**Figs. 4d-4k**). These results are consistent with the protective effects observed with the CD11b activator ONT01, mechanistically confirming that CD11b activation regulates TLR7-driven inflammatory pathways, systemic autoimmunity, and kidney damage.

### Oral CD11b activator is therapeutic in a genetic model of LN

The MRL/lpr mice that develop high circulating levels of IFN-I, systemic inflammation and multi-organ pathology, including skin and kidney damage serve as a reliable genetic model of LN [70]. We previously showed that CD11b activation ameliorates SLE in this model system [41]. We hypothesized that age-dependent increase in albuminuria in this model is driven by elevated serum suPAR levels and that CD11b activation could suppress it. Vehicle-treated MRL/lpr mice exhibited whisker loss, alopecia, and multiple skin lesions (**Figs. 5a-5d**). They also had significantly increased spleen weight and size, and splenic expansion of CD11b+ leukocytes (red bars, **Figs. 5e-5g**). Circulating levels of anti-dsDNA antibodies, suPAR, and IL-6 were significantly higher compared to haplotype-matched MRL/mpj controls (gray vs. red bars, **Figs. 5h-5j**). Expectedly, vehicle-treated mice showed significantly higher albuminuria (**Fig. 5k**), significant leukocyte glomerular infiltration, glomerular damage, and increased leukocyte infiltration in the thickened epidermis (**Figs. 5l-5m**). IF of kidney glomeruli showed increased IgG and C3 deposition and decreased Synpo and active integrin β1 staining (**Figs. 5n-5o**). Oral ONT01 significantly reduced alopecia, skin lesions, normalized spleen size and weight, splenic CD11b+ leukocyte expansion, and epidermal thickening (dark blue bars, **Figs. 5b-5o**). It also significantly reduced circulating levels of anti-dsDNA antibodies, suPAR, and IL-6, albuminuria, leukocyte infiltration, glomerular damage, and podocyte loss, mimicking the protective effects observed in TLR7-dependent models.

**Figure 5.**
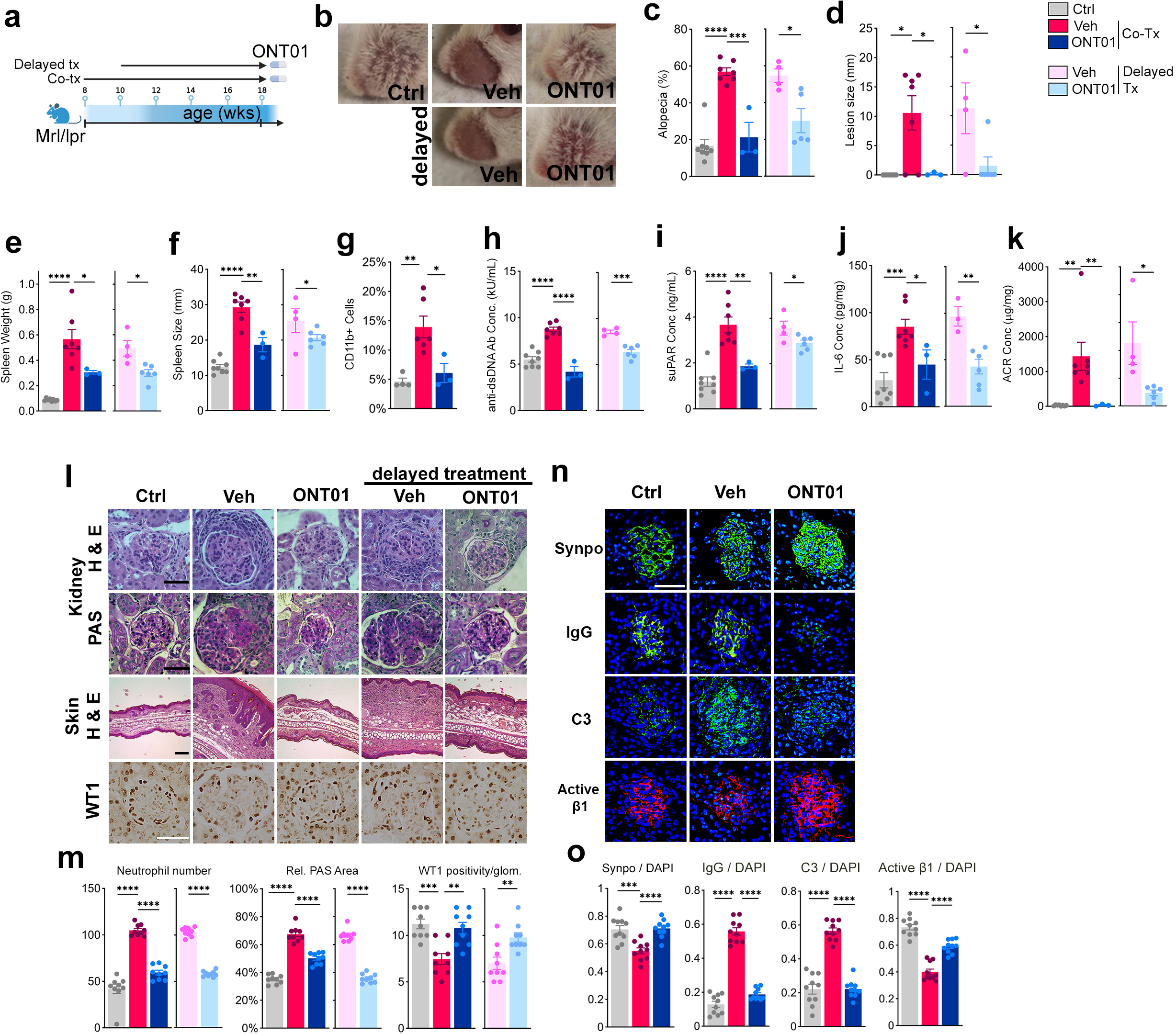
ONT01 protects against LN in a genetic mouse model. **(a)** Experimental diagram showing plan of *in vivo* treatment of MRL/*lpr* genetic model of LN with ONT01. MRL/*MpJ* were used as control. Arrows indicate the duration of treatment with ONT01. Treatment was initiated either at the time of disease onset (when animals were 8-wk old) or 2-weeks after the disease onset (when the animals were 10-wk old). **(b)** Representative images showing facial alopecia from various groups of animals: MRL/*MpJ* control (Ctrl), MRL/*lpr* treated with vehicle alone starting at 8-wks of age (Veh), MRL/*lpr* treated with ONT01 starting at 8-wks of age (Onteg.), MRL/*lpr* treated with vehicle alone starting at 10-wks of age (Veh delayed), MRL/*lpr* treated with ONT01 starting at 10-wks of age (Onteg. delayed). **(c-d)** Graphs showing quantification of relative level of alopecia (**c**) and skin lesion size (in mm) (**d**) at the end point from various groups of animals, as in **b**: MRL/*MpJ* control (Ctrl, gray bars), MRL/*lpr* treated with vehicle alone starting at 8-wks of age (Veh, red bars), MRL/*lpr* treated with ONT01 starting at 8-wks of age (Onteg., dark blue bars), MRL/*lpr* treated with vehicle alone starting at 10-wks of age (Veh delayed tx purple bars), MRL/*lpr* treated with ONT01 starting at 10-wks of age (Onteg. delayed tx, light blue bars). Each bar shows mean ± SEM (control n=8, vehicle n=7, ONT01 n=3, vehicle delayed n=4, and ONT01 delayed n=6 animals/group). *p<0.05, ***p<0.001. **(e-f)** Graphs showing quantification of spleen weight (in g) and size (in mm) at the end point from various groups of animals, as in **b-d**. *p<0.05, **p<0.01, ****p<0.0001. **(g)** Graph showing flow cytometric analysis of CD11b+ splenocytes at week 18 (end point) from Ctrl, Veh. and Onteg. groups of animals. Each bar shows mean ± SEM. *p<0.05, **p<0.01. **(h-j)** Graphs showing ELISA based quantification of anti-dsDNA antibody, suPAR and IL-6 levels in the end-point sera from various groups of animals, as in **b**. Each bar shows mean ± SEM. *p<0.05, **p<0.01, ***p<0.001, ****p<0.0001. **(k)** Graph showing albumin to creatinine ratio (ACR) in the end-point urine from various groups of animals, as in **b**. Each bar shows mean ± SEM. *p<0.05, **p<0.01. **(l)** Representative images from end point kidney (stained with H&E and PAS) and skin (stained with H&E) sections from various groups of animals, as in **b.** Also presented are representative images of kidney glomeruli from immunohistochemical staining of kidney with antibodies against Wilms Tumor 1 (WT1, brown) from various groups of animals, as in **b**. Scale bar: 50 μm for kidney; 100 μm for skin. **(m)** Bar graphs showing quantification of leukocyte numbers, relative PAS area, and WT1 positive cell numbers among the experimental groups presented in **l**. Each bar shows mean ± SEM (n=8-10 independent images/group). **p<0.01, ****p<0.0001. **(n)** Representative immunofluorescence images of kidney glomeruli from staining of kidney with antibodies against synaptopodin (Synpo, green), IgG (IgG, green), complement (C3, green) and integrin β1 activation epitope (9EG7, red) from animals as in **l**. Nuclei were stained with DAPI (blue). Scale bar: 50 μm. **(o)** Bar graphs showing quantified ratios of IgG and C3 deposition in glomerulus of the various experimental groups presented in **n**. Each bar shows mean ± SEM (n=8-10 independent images/group). ****p<0.0001.

To evaluate the therapeutic potential of ONT01, we delayed ONT01 administration by two weeks and initiated MRL/lpr mice treatment at 10 weeks of age (**Fig. 5a**). Delayed ONT01 treatment significantly reduced alopecia and skin lesions (pink vs. light blue bars, **Figs. 5b-5d**), normalized spleen size, weight, splenic CD11b+ leukocyte numbers, and epidermal thickening, and significantly reduced circulating levels of anti-dsDNA antibodies, suPAR, IL-6, albuminuria, leukocyte infiltration, glomerular damage, and podocyte loss (pink vs. light blue bars, **Figs. 5b-5o**). These results further support ONT01 as a novel LN therapeutic.

### ONT01 treatment in a humanized mouse model of LN provides therapeutic benefit

To examine the role of CD11b activation in human context, we developed a murine model engrafted with human PBMCs using our recently described technique (**Fig. 6a**) [7]. NSG mice were injected intravenously with PBMCs from healthy donors or LN patients. Engraftment rates were confirmed by quantifying human CD45+ cells in the blood, bone marrow (BM) and spleen of the recipient mice (**Fig. S7a-S7e**). Analysis of the patient sera showed significantly elevated suPAR and anti-dsDNA antibody levels, as compared with healthy controls (**Fig. S7f-S7g**). Mice engrafted with patient PBMCs developed lupus like pathology, prominent alopecia, systemic inflammation and kidney damage (**Figs. 6a-6c**), and showed significant splenic expansion of CD11b+ leukocytes (**Figs. 6d** and **S7h-S7i**), splenomegaly (**Fig. S7j**), increased anti-dsDNA antibodies, suPAR, IL-6 and IFNβ in serum (**Figs. 6e-h**), and increased albuminuria (**Fig. 6i**). Histological analyses showed significant glomerular damage and leukocyte infiltration in kidney sections (**Figs. 6j-6m**).

**Figure 6.**
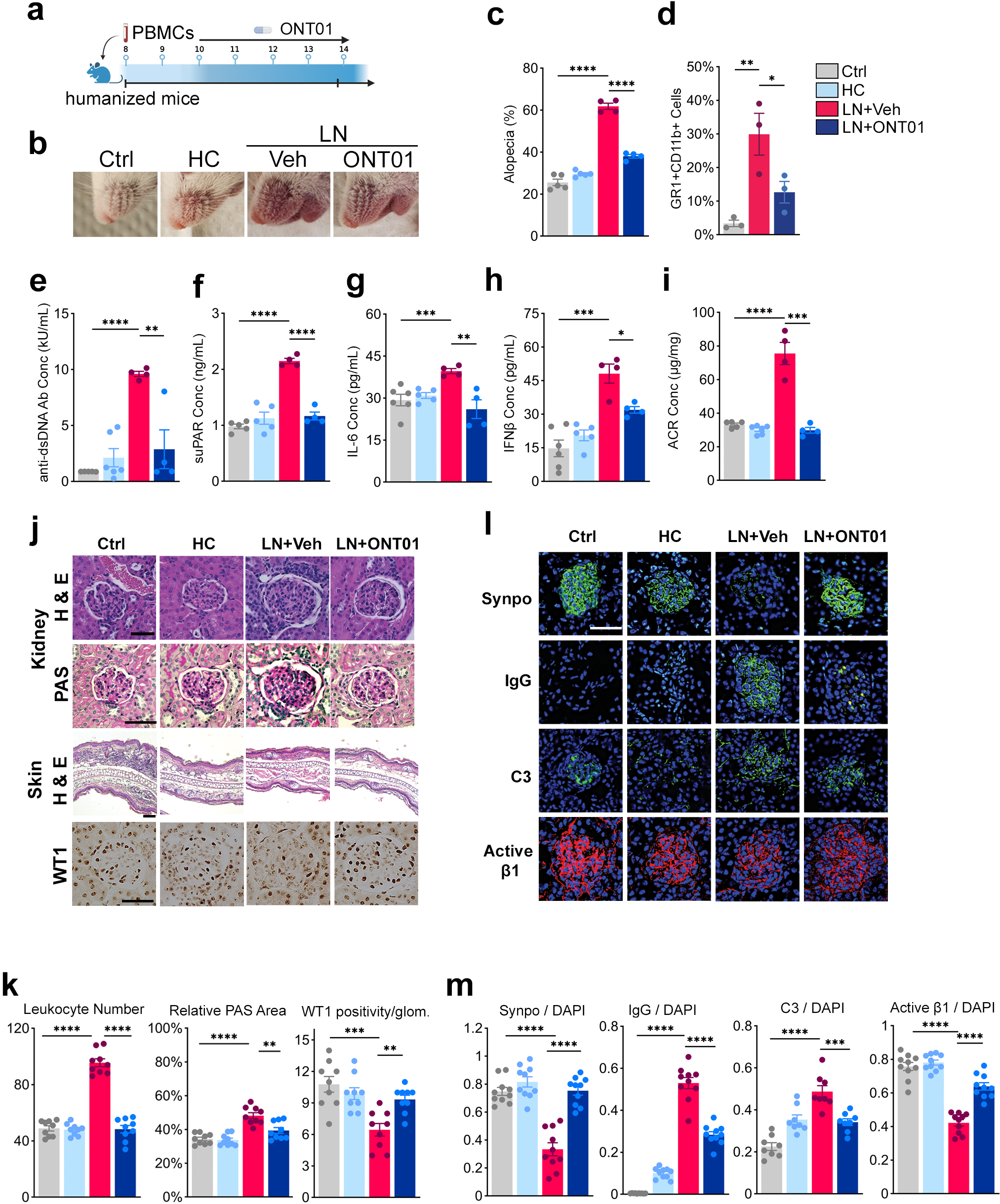
Transfer of LN PBMCs recapitulates human like disease in mice that can be suppressed with CD11b activator ONT01. **(a)** Experimental diagram showing plan of human PBMC transfer and engraftment into NOD-scidIL2γnull NSG mice to generate a humanized LN model and its treatment with ONT01. Arrow indicates the time of intiation of treatment with ONT01 (10-wk of age), two weeks after transfer of PBMCs. **(b)** Representative images showing facial alopecia from various groups of animals: Non-transferred control (Ctrl), healthy PBMC transfer control (HC), animals with LN PBMC and treated with vehicle alone (Veh) and animals with LN PBMC and treated with ONT01 (Onteg.). **(c)** Graph showing quantification of relative level of alopecia at the end point from various groups of animals, as in **b**: non transferred control (Ctrl, gray bars), Healthy PBMC transferred control (HC, light blue bars), LN PBMC transferred mice and treated with vehicle alone (Veh, red bars), and LN PBMC transferred mice and treated with ONT01 (Onteg., dark blue bars). Each bar shows mean ± SEM (control n=5, HC n=5, vehicle n=4, ONT01 n=4 animals/group). ****p<0.0001. **(d)** Graph showing flow cytometric quantification of GR1+CD11b+ splenocytes at end point from Ctrl, Veh. and Onteg. groups of animals. Each bar shows mean ± SEM. *p<0.05, **p<0.01. **(e-h)** Graphs showing ELISA based quantification of anti-dsDNA antibody, suPAR, IL-6 and IFNβ levels in the end-point sera from various groups of animals, as in **b**. Each bar shows mean ± SEM. **p<0.01, ***p<0.001,****p<0.0001. **(i)** Graph showing albumin to creatinine ratio (ACR) in the end-point urine from humanized mice treated with either vehicle or ONT01, as in **b**. Each bar shows mean ± SEM (n=5/group). **p<0.01, ***p<0.001. **(j)** Representative images from end point kidney (stained with H&E and PAS) and skin (stained with H&E) sections from various groups of animals, as in **b.** Also presented are representative images of kidney glomeruli from immunohistochemical staining of kidney with antibodies against Wilms Tumor 1 (WT1, brown) from various groups of animals, as in **b**. Scale bar: 50 μm for kidney; 100 μm for skin. **(k)** Bar graphs showing quantification of leukocyte numbers, relative PAS area, and WT1 positive cell numbers among the experimental groups presented in **j**. Each bar shows mean ± SEM (n=8-10 independent images/group). **p<0.01, ***p<0.001, ****p<0.0001. **(l)** Representative immunofluorescence images of kidney glomeruli from staining of kidney with antibodies against synaptopodin (Synpo, green), IgG (IgG, green), complement (C3, green) and integrin β1 activation epitope (9EG7, red) from animals as in **j**. Nuclei were stained with DAPI (blue). Scale bar: 50 μm. **(m)** Bar graphs showing quantified ratios of IgG and C3 deposition in glomerulus of the various experimental groups presented in **l**. Each bar shows mean ± SEM (n=8-10 independent images/group). ****p<0.0001.

ONT01 treatment of LN PBMC-engrafted mice significantly reduced systemic autoimmunity and kidney damage. It reduced alopecia, normalized splenic CD11b+ leukocyte levels, decreased serum anti-dsDNA antibodies, suPAR, IL-6, and IFNβ levels, and normalized albuminuria (**Figs. 6b-6i**). IHC showed reduced leukocyte infiltration and kidney damage in ONT01 treated mice (**Figs. 6j-6k**) and IF confirmed reduced IgG and C3 immune complex deposition, and improved Synpo and active integrin β1 staining (**Figs. 6l-6m**).

Collectively, these data show that engrafting human LN PBMCs into immunodeficient mice successfully recapitulates human LN features, providing a useful model for studying LN. Our results also demonstrate that CD11b activation with ONT01 reduces LN-induced systemic inflammation and kidney damage, highlighting its therapeutic potential.

## Discussion

Our study examined the association of *ITGAM* coding variants with lupus nephritis (LN) and the functional role of CD11b, encoded by *ITGAM*, in disease pathogenesis and therapeutics. We found that coding variants in *ITGAM* correlate with elevated serum suPAR levels, a known inflammatory cytokine and kidney disease biomarker, suggesting a functional link. Importantly, pharmacologic activation of CD11b by ONT01 suppresses suPAR production and TLR7-dependent signaling in myeloid cells, ameliorating LN in multiple murine models, including a humanized model with LN patient PBMCs. These findings provide new insights into the molecular mechanisms linking *ITGAM* SNPs, suPAR, and TLR7 signaling in LN and support the development of CD11b activator ONT01 as a novel, mechanism-based therapeutic for LN.

Integrin CD11b plays a critical role in regulating immune responses, particularly in the adhesion, migration, and signaling functions of leukocytes. Previous studies have shown that *ITGAM* SNPs increase the risk and severity of LN and other kidney dysfunctions [26–28, 31, 46–49]. Studies also showed that these SNPs affect the expression and function of CD11b on the surface of leukocytes [35–38, 71]. However, how these SNPs contribute to LN pathogenesis has not been completely clear. Our study shows that absence of CD11b increases the production of suPAR by myeloid cells, and that *ITGAM* SNPs are associated with elevated serum suPAR levels in LN patients, suggesting that *ITGAM* variants contribute to LN pathogenesis by increasing myeloid-derived suPAR production.

We also discovered that pharmacologic or genetic gain-of-function mutation mediated activation of CD11b can suppress TLR7-dependent signaling in myeloid cells. TLR7 is a key driver of SLE and LN [15–17, 19, 20, 72]. TLRs are aberrantly activated via binding to self-ligands, that recruits adaptors MyD88 and TRIF to the cell membrane to assemble myddosome signaling complex [52, 63, 73]. The myddosome complex rapidly dissociates from the cell membrane to assemble additional adaptors, kinases and effectors to launch a signaling cascade [63]. This results in activation of transcription factors, such as NF-κB, IRF3 and IRF7, to increase IFN I and other pro-inflammatory cytokines, such as IL-6 and TNF-α [52]. These subsequently promote autoantibody production, immune complex formation, and tissue inflammation [15, 17, 19, 20, 74]. Gain-of-function mutations in TLR7 result in its hyperactivation, amplify the immune response to self-antigens, and are associated with increased risk of SLE and LN [20]. Thus, several TLR7 antagonists and antibodies are in pre-clinical and clinical development and show promising results in early SLE trials, though data in LN with these is limited [59, 75–78]. However, global or cell-specific genetic deletion of TLR7 provides only modest reductions in kidney disease in SLE models [79], perhaps because the myddosome rapidly dissociates from the cell membrane [63]. Here, our findings suggest CD11b activation is a novel strategy to regulate TLR-dependent inflammatory pathways. We found that TLR7 activation increases production of suPAR, IFN-β, IL-6, and TNF-α, in line with past literature [7–9, 39, 52, 63], and that CD11b activation with ONT01 suppresses these inflammatory mediators. We found that this suppression is mediated via suppression of the NF-κB and IRF3/7 pathways. CD11b activation attenuated key myddosome components MyD88 and TRIF and regulated the phosphorylation of key signaling components pSrc, MyD88, TRIF, pTBK1, pSTING, and NLRP3, downstream of TLR7 activation. These findings identify a key role for CD11b in regulating TLR7-dependent immune responses in myeloid cells and suggest CD11b activation as a novel strategy to regulate TLR-dependent inflammatory pathways.

In multiple experimental models, we demonstrated that CD11b activation has therapeutic potential in LN. ONT01 treatment reduced systemic autoimmunity and kidney damage, evidenced by decreased alopecia, splenomegaly, serum anti-dsDNA antibodies, serum suPAR, IL-6, IFN-β, and TNFα levels, and albuminuria and glomerular damage. Healthy glomerular podocytes and their retention in the kidney is essential for normal kidney function. Podocyte attachment to the glomerular basement membrane retains them in a healthy kidney and is mediated via its surface expressed integrin β1. The levels of active β1 and the podocyte-specific marker synaptopodin (Synpo) correlate with kidney function [80, 81]. ONT01 significantly reduced the loss in glomerular podocytes and protected podocyte expressed synaptopodin and active integrin β1 levels. These protective effects were consistent across different murine models, reinforcing the therapeutic efficacy of CD11b activation. Our study also confirms the mechanism of action of CD11b activation by using a genetic CD11b knock-in mouse model that expresses a gain-of-function active mutant of CD11b, mimicking CD11b pharmacologic activation [44]. We found that genetic CD11b activation also protected against TLR7-dependent LN, while absence of CD11b worsened the disease, consistent with prior studies showing CD11b deficiency to be pathogenic in SLE [69]. These results are also consistent with previous studies showing the therapeutic efficacy and safety of CD11b activation by ONT01 in other models of inflammatory and autoimmune diseases, such as SLE, cardiovascular injury and acute and chronic kidney injury [42, 56, 58, 62, 82–87]. These data suggest that CD11b activation is a therapeutic mechanism to suppress TLR7-dependent autoimmune disease. Moreover, in a recently complete human Phase 1 clinical trial (KEYNOTE A-36), ONT01 was found to be well tolerated [55, 85, 88, 89], suggesting its safety for future clinical studies in LN.

Collectively, our findings identify CD11b as an important regulator of TLR7-dependent inflammatory diseases and strongly support the development of CD11b agonists as novel therapeutics for LN and other autoimmune diseases. We expect that future studies will explore the clinical application of CD11b activation in human patients and would pave the way for innovative treatments targeting innate immune cell regulation in autoimmunity.

## Methods

### Reagents and antibodies

ONT01 was a gift from Allosite Therapeutics (Miami, FL). Bovine serum albumin (BSA) was from Sigma (St. Louis, MI). Non-fat milk was obtained from BioRad (Hercules, CA). PCR reagents were obtained from New England Biolabs Inc (Beverly, MA). All cell culture reagents were purchased from Thermo Fisher Corp (San Diego, CA). LPS (cat# tlrl-eklps), Pam2 (cat# tlrl-pm2s-1), Pam3 (cat# tlrl-pms), PMA (cat# tlrl-pma), polyI:C (cat# tlrl-plc) and CpG (cat# tlrl-2216) were purchased from Invivogen (San Diego, CA). Human and mouse recombinant cytokines and ELISA kits were purchased from R&D Systems and were declared by the manufacturer to contain <0.1 ng of LPS per μg of protein. The following primary antibodies for western blot were purchased from Cell Signaling Technologies (Danvers, MA): Phospho IKK-α/β (cat# 2078), IKK-β (cat# 8943), Phospho p65/NF-kB (cat# 3036), MyD88 (cat# 4283), GAPDH (cat# 2118), Phospho IRF3 (cat# 4947), IRF3 (cat# 4302), Phospho IRF7 (cat# 14767), Phospho c-Cbl (cat# 8869), c-Cbl (cat# 8447), Phospho Src (cat# 6943), and Src (cat# 2108). Other antibodies used were anti-rabbit IgG-HRP conjugate (cat# W4011, Promega, Madison, WI), anti-mouse IgG-HRP conjugate (cat# W4021, Promega, Madison, WI), NF-kB (cat# 16502, Abcam, Cambridge, MA), and IRF7 (cat# PA5-20280, Thermo Scientific, Waltham, MA). Antibodies used for immunofluorescence experiments were as follows: WT1 (cat# SC-192, Santa Cruz), synaptopodin (cat# SC-21537, Santa Cruz), activated β1 (cat# 553715, BD Biosciences) C3 (cat# GC3-90F-Z, ICL, Inc.) or IgG (cat# A11029, Thermofisher Scientific). The following antibodies for flow cytometry experiments were purchased from Biolegend (San Diego, CA): APC/Cyanine7 anti-mouse CD45 (cat# 103115), PE anti-mouse Ly-6G (cat# 127607), APC anti-mouse Ly-6C (cat# 128015), FITC anti-mouse Ly-6G/Ly-6C (Gr-1) (cat# 108406), APC anti-mouse/human CD11b (cat# 101212), APC/Cyanine7 anti-mouse CD45 (cat# 103116), Alexa Fluor 488 anti-mouse/human CD11b (cat# 101217). The following antibodies for flow cytometry were purchased from BD Biosciences: BD Pharmingen FITC Rat Anti-Mouse Ly-6G and LY-6C (cat# 553127), BD Horizon PE-CF594 Rat Anti-CD11b Antibody (cat# 562287). LIVE/DEAD Fixable Aqua Dead Cell Stain Kit was purchased from Thermofisher Scientific (cat# L34966).

### Human samples

Data from 131 active lupus cases consisting of patients of self-reported European ancestry that had serum available for suPAR analysis were obtained from Rush University Medical Center and Mayo Clinic. Informed consent was obtained from all patients in all the cohorts included in this study, and the study was approved by the Institutional review boards at the respective institutions.

### Genotyping

Patients were genotyped as described previously (33). Briefly, genotyping at coding change SNPs in *ITGAM* (rs1143678, rs1143679 and rs1143683) was performed using custom designed Applied Biosystems Taqman primers and probes on an Applied Biosystems 7900HT PCR machine with >98% genotyping success. Genotyping scatter plots were all reviewed individually for quality, and genotype frequencies did not deviate significantly from the expected Hardy-Weinberg proportions (p>0.01).

### Human PBMC isolation

Human peripheral blood mononuclear cells (PBMCs) were isolated from fresh blood collected from healthy volunteers under an IRB-approved protocol and using RosetteSep (human monocyte Enrichment Cocktail, Cat# 15068 Stemcell Technologies, Vancouver, Canada) followed by Ficoll-Hypaque density gradient centrifugation according to published protocols [90]. Briefly, blood was diluted with PBS containing 2% FBS and was layered on the top of the ficoll-paque solution very gently. The tubes were centrifuged at 400g for 20 minutes at room temperature using slow acceleration and deceleration, and the upper plasma layer was removed carefully. The buffy coat (PBMCs) from plasma-ficoll interphase was transferred to a clean tube and DPBS was added and mixed well. Again, it was spun at 100g for 10 minutes to remove any contaminating Ficoll and platelets/proteins. The cells were washed with DPBS, resuspended in RPMI-1640 with GlutaMAX (Cat# 61870-036, GIBCO), supplemented with 10% FBS and seeded into T75 (75 cm^2^) tissue culture flask (Cat#156800, Thermo). Non-adherent cells were removed by gentle pipette aspiration after 2 h of incubation at 37°C in a humidified atmosphere containing 5% CO2. Subsequently, an equal volume of fresh complete medium was added to each flask and the cells were placed back in the incubator for approx. 24h [91, 92].

### Cell line

The cell line RAW 264.7 murine macrophage was obtained from the American Type Culture Collection (ATCC #TIB-71) and was maintained accordingly.

### Mice

The C57BL/6J (B6) wild type (cat# 000664) (CD11b WT), B6 CD11b^-/-^ (cat# 003991) [93] (CD11b KO), female MRL/lpr (cat# 000485) and haplotype-, age- and sex-matched control MRL/MpJ (cat# 000486) were from The Jackson Laboratory, Bar Harbor, ME. To examine role of genetic activation of CD11b on TLR7-dependent lupus nephritis, we used transgenic mice that we recently generated expressing a constitutively activated CD11b by introducing an I332G point mutation in the murine Itgam gene (C57BL/6 *ITGAM*^I332G^) [44] (CD11b KI). The mice were maintained in specific-pathogen-free conditions and used in accordance with the Institutional Animal Care and Use Committee (IACUC) and the respective institutional guidelines. The MRL/MpJ mice were used as a non-lupus prone, haplotype-matched strain to compare lupus-prone mice to normal pathology.

### TLR7-induced mouse model

8-10 weeks old C57BL/6J wild-type, CD11b^-/-^ or CD11b KI female mice were treated with imiquimod (IMQ) according to published protocols to induce TLR7-induced systemic lupus [43]. Briefly, the skin on the right ears of the mice were treated topically with 1.25 mg of 5% imiquimod cream three times per week for 8-10 weeks. The ultraviolet B (UVB) irradiation was also applied for 30s using Dermaray M-DMR-1 lamp bulbs with peak emission at 311 nm. For the treatment groups, mice were treated with ONT01 (30mg/kg BID) or vehicle (1% Tween-20 in sterile saline) orally twice daily. Urine was collected every other week. At the end of the experiment, blood was collected by terminal cardiac puncture and processed for sera and PBMCs. Ears and kidneys were harvested for RNA, protein, and histology. Macrophages were generated from bone-marrow derived monocytes and cultured in RPMI containing 10% FBS cultured for 5-7 days. Spleen was isolated for measurement of size and weight and for flow cytometric analyses.

### MRL/lpr mouse model

8-weeks old mice were obtained from the Jackson Laboratory (Bar Harbor, ME). Female MRL/lpr mice were treated orally with ONT01 (30mg/kg/twice a day) or vehicle (1% Tween-20 in sterile saline) twice daily, beginning at 8-10 weeks of age until euthanasia at 20 weeks of age (*19*). Haplotype-matched female MRL/MpJ mice were used as phenotypic controls. Kidneys and affected skin samples were harvested after perfusion with PBS (Phosphate Buffered Saline). Regions of alopecia were measured on the face and dorsum, with greatest diameter determined to the nearest millimeter with calipers. Paraffin embedded sections (4 μm) were stained with hemotoxylin-eosin (H&E) or periodic acid-Schiff (PAS). The assessments were blinded for histological analysis. Sample size was chosen based on previous studies using interventions in MRL/*lpr* mice to assess similar outcomes in phenotype [94].

### Humanized mouse model

Wild-type female C57BL/6J mice or NOD-scidIL2γnull NSG mice were purchased from the Jackson Laboratories. 6–8 weeks old mice were housed under pathogen-free conditions in the animal facility at the Rush University Medical Center in accordance with the institutional guidelines under Institutional Animal Care and Use Committee approval. Human LN patient blood was used and isolated PBMCs were injected intravenously (i.v.) into NSG mice (2.5 x 10^6^ cells per mouse) on day 0. The humanized mice were monitored daily for significant clinical signs (alopecia, skin lesions, weight loss & arthritis), morbidity and mortality. Body weights were recorded weekly and skin lesions were photographed and scored weekly.

### Whole blood sampling for serum and FACS analysis of human cell engraftment rates

A small sample of peripheral blood was drawn from the mice under general anesthesia 2-, 4- and 6-weeks post-humanization by penetrating the retro-orbital sinus in mice with a glass capillary tube. The blood sample was split for FACs analysis and serum preparation. For FACs staining, an anticoagulant was added to the blood, RBCs were lysed, and the isolated leukocytes were stained with fluorescence-labeled antibodies against human CD45 and mouse CD45 for flow cytometry-based quantification to confirm high human cell engraftment rates. For serum, the whole blood was clotted at room temperature for 30 minutes, followed by a 10-minute centrifugation at 2,000g. The RBC free serum was transferred into micro-centrifuge tubes and stored at −80° C until further analysis.

### Preparation of murine bone marrow derived macrophages

Mice were euthanized via CO2 and secondary cervical dislocation, and the bone marrow cells from tibia and femurs were collected by flushing with DMEM containing 10% FBS and using a 25-gauge needle according to published protocols [41]. Any cell clumps were dispersed by gently pipetting the solution and then passed through a 70μm cell strainer. The suspension was subsequently centrifuged, the supernatant discarded, and the pelleted cells gently resuspended in DMEM containing 10% FBS. Red blood cells were removed by treating the cell suspension with RBC lysis buffer according to manufacturer’s instructions. Cells were washed and resuspended in DMEM containing 10% FBS. For generation of bone marrow derived macrophages, approx. 8X10^6^ cells were seeded in 6 mL of DMEM containing 10% FBS and 50 ng/mL MCSF (PeproTech #315-02) in a 100mm plate on day 0. Cells were cultured for 7 days and fed with an additional 3mL media with 50ng/mL MCSF on day 4 of culture. On day 7, cells were removed from the plates with an EDTA solution, viability was checked, and cells were replated using DMEM with 1% FBS before planned treatments.

### NF-κB and STING Agonist treatment

CD11b KI BMDMs were treated with NF-κB agonist (Phorbol myristate acetate, PMA) at 200 ng/mL and STING agonist (ADU-S100) at 1μg/mL for 4 hours. Cell supernatant was used in ELISA experiments.

### Analysis of albuminuria

Mouse urine samples were collected, transferred into a microcentrifuge tube and stored at −80° for future analysis. Urinary albumin concentration was measured using a mouse albumin ELISA (Bethyl Laboratories, Montgomery, TX #E99-134) and urinary creatinine assay (Exocell, Philadelphia, PA #1012), respectively, according to manufacturer’s protocol. Subsequently, urine albumin:creatinine (ACR) ratio was calculated.

### Quantification of suPAR, and anti-dsDNA Antibody

Mice sera and RAW cell supernatants were used for quantification. Commercial ELISA kits for murine anti-dsDNA antibodies (Alpha Diagnostic #5120), and suPAR (R & D system #DY531) were performed according to the manufacturer’s instruction.

### ELISA assays for cytokine readout

Cell supernatant and blood were collected for cytokine analysis by ELISA and were prepared by centrifugation and isolation of supernatant which was frozen until use. Cytokine levels (IFN-b, IL-6, TNF-a) were measured by ELISAs ordered from R&D Systems and were performed according to manufacturer’s protocol. Briefly, mouse macrophages were cultured in the presence of vehicle (1% DMSO), ONT01 (20 µM), IMQ (20µg/mL), or IMQ + ONT01 for 6 h and the cell supernatant was collected. Mouse cell culture supernatants were assayed using commercially available sandwich ELISA kits for mouse IL6 (R&D Systems, Minneapolis, MN, catalog# M6000B), and TNF-α (R&D Systems, Minneapolis, MN, catalog# MTA00B) according to manufacturer’s instruction. For measuring IFN-β levels, mouse macrophages (WT and CD11b^-/-^ and KI) were cultured in the presence of vehicle (1% DMSO), ONT01 (20 µM), IMQ (20µg/mL), or IMQ + ONT01 for 12h and the cell culture supernatant were assayed using commercially available sandwich ELISA kits for mouse IFN-β (PBL Assay Science, Piscataway, NJ, catalog# 42400-1) according to manufacturer provided instructions.

### Histology

One part of the removed kidney was fixed in 10% formalin and embedded in paraffin and another part was immediately snap frozen in OCT on liquid nitrogen and stored at −80°C. Samples were labelled and sent to the University of Illinois at Chicago Research Histology Core for sectioning and staining. Paraffin embedded sections (4 μm) were stained with hemotoxylin-eosin (H&E) or periodic acid-schiff (PAS). Stained slides were blindly evaluated by an experienced pathologist for chronic tubulointerstitial damage, glomerular sclerosis, and tubulointerstitial inflammation using 0 (none), 1+ (<25%), 2+ (25-50%), 3+ (>50%) scoring system. Formalin-fixed and frozen sections of the skin were prepared as for kidneys.

### Immunofluorescence

For immunofluorescence studies, tissue sections (4 μm) were cut and fixed in −20°C acetone before immunofluorescence staining. Sections were blocked (4% FBS, 4% BSA, 0.4% Fish gelatin in PBS) at room temperature for 1 hr and incubated with the appropriate primary antibodies in blocking buffer at 4°C overnight. Sections were incubated with the appropriate secondary antibody (Thermofisher Scientific) and mounted with DAPI medium (Vector Laboratories). Fluorescence images were acquired using a Zeiss LSM 700 confocal microscope with a PLAN-Apochromat 20 X objective and an AxioCam camera and analyzed using the Zen software (Carl Zeiss Group). The number of WT1 positive cells per glomeruli was calculated manually using the confocal microscope (5 glomeruli/tissue). The fluorescence intensities for synaptopodin, C3, 9EG7 and IgG were calculated using the ImageJ Software (5-8 glomeruli/tissue).

### Flow cytometry of mouse splenocytes for immunophenotyping

Single cell suspensions of mouse splenocytes were prepared using a cell strainer, rinsing with FACS buffer. The cells were washed with PBS and then underwent RBC lysis using 1x multi-species RBC Lysis Buffer (Cat#00-4333-57, eBioscience). The cells were washed again in PBS and resuspended in FACS buffer (2% FBS in PBS) at 1.0 x 10^6^ cells/mL for analysis by flow cytometry. After blocking for 15 minutes, cells were resuspended in 100uL FACS buffer and incubated with 2uL each of respective antibodies or isotype control for 30 minutes/ 1 hour on ice, fixed with BD stabilizing fixative, quantified by flow cytometry using LSR Fortessa research only use (ROU) or BD FACSCanto RUO and data analyzed using FlowJo software (FlowJo LLC). Cutoff values for positive staining were determined using compensation controls for each fluorophore. Results were reported as number of cell subsets/million splenocytes.

### Quantification of inflammatory markers by qPCR

RAW 264.7 cells in culture were washed with PBS and lysed using 700μL of <u>Trizol</u>. Kidney tissues were homogenised in Trizol followed by RNA extraction using RNeasy Mini kit (catalog #74106, Qiagen) according to manufacturer’s protocol. cDNA was synthesized using iScript Reverse Transcription Supermix (cat #1708841, BioRad) according to manufacturer’s instruction. cDNA, desired primers, and SsoAdvanced Universal SYBR Green Supermix (cat #1725271, BioRad) were subjected to quantitative Real-time PCR using thermal cycler according to the manufacturer’s instruction and relative changes in gene expression were analyzed using the formula 2^-ΔΔCt^. All inflammatory marker genes including housekeeping gene (PPIA/GAPDH) were quantified. The primer sequences for the uPAR, IFNβ, IL6, and TNFα genes have been described by our group previously [41].

### Western Blot

Cells were lysed with RIPA buffer containing EDTA (Cat #BP-115D, Boston Bioproducts) and cocktails of protease inhibitor (Catalog #11836170001, Millipore Sigma) and phosphatase inhibitor (cat #PI78420, Thermo Fisher Scientific). Protein concentration was calculated with the Pierce BCA Protein Assay Kit (Cat #23227, Thermo Fisher Scientific) according to manufacturer’s protocol and the samples were aliquoted accordingly before freezing for future use. Samples were thawed on ice. NuPAGE LDS Sample Buffer (catalog #NP0007, Invitrogen) with 1% β-mercaptoethanol was added to each sample before denaturing them at 90°C for 5 minutes. Equal amounts of proteins were loaded in wells of a 4-12% NuPAGE Bis-Tris Gel (cat #NP0322BOX, Invitrogen) and blotted to a PVDF membrane (catalog #IPFL00010, EMD Millipore). Subsequently, the membrane was blocked with blocking buffer (5% BSA in PBS) for 1 hour at RT followed by incubation with primary antibodies in assay buffer (0.5% BSA and 0.05% Tween-20 in PBS) overnight at 4°C. Subsequently, the membranes were washed with assay buffer and incubated with anti-rabbit or anti-mouse secondary antibody in assay buffer for 1 hour at RT. Proteins were detected using either a LI-COR odyssey imager or incubation with Pierce ECL western blotting substrate and imaged with X-ray film. Data presented is representative of at least three independent experiments.

### Statistical analysis

To calculate statistical significance, a two-tailed student’s t-test was used, unless otherwise specified. The differences in cytokine array concentrations were statistically analyzed using the Mann-Whitney test and plotted with GraphPad Prism software package. An unpaired t-test was used for comparison between two independent groups. Welsch’s t-test was used for comparison between two populations with equal means. For multiple comparisons, one-way ANOVA with Tukey’s test was used. A value of p<0.05 was considered statistically significant. Unless otherwise specified, results are represented as mean ± SEM.

### Study approval

The studies on human samples were reviewed and approved by Institutional review boards at the Rush University Medical Center, Mayo Clinic and Hospital for Special Surgery. All subjects provided appropriate prior informed consent to participate in the study. The murine studies were reviewed and approved by the Institutional Animal Care and Use Committee (IACUC) at Rush University Medical Center.

## Acknowledgements

We thank current and past members of the Gupta laboratory for their technical help and scientific discussions on this long-running project. We also thank Roberto Vazquez-Padron for the KI animals. This work was supported in part by grants R01DK084195, R01DK136297, R01 CA244938 to VG from the National Institutes of Health (NIH), grant MJFF-022480 from Michael J Fox Foundation to VG, a predoctoral fellowship from the NIH to VV (F31DK129006), a grant from NIH to DL-R (R43AI179510) and with resources from the Rush University Medical Center (RUMC) and from the University of Texas Medical Branch (UTMB).

## Author Contributions

XL, VJ, BN, SQK, VV, HF designed and performed *in vitro* and *in vivo* experiments; NRC, KA, HW, DCW, DL-R, SBP, AB contributed to *in vitro* experiments and data analyses; VV, KA, MJ, JMD and MAJ contributed to human samples and analyses; XL, SQK, BN and DJC performed scoring of lupus murine skin and lupus murine kidneys; RIVP provided CD11b KI model and expertise; MJ, JR, TN provided disease expertise; HF, TN and VG co-designed and co-supervised the study and co-wrote the paper.

